# Reversal of obesogenic feeding and hypometabolism by a bifurcating GABAergic neural circuit

**DOI:** 10.1101/2022.01.23.477433

**Authors:** Yong Han, Yang He, Lauren Harris, Yong Xu, Qi Wu

**Affiliations:** USDA/ARS Children’s Nutrition Research Center, Department of Pediatrics, Baylor College of Medicine, Houston, TX, USA

## Abstract

Diet-induced obesity is characterized by unsatiated consumption of energy-dense diets and impaired metabolism, whereby anti-obesity effect of the high-level of circulating leptin is unknowingly blunted. Emerging evidence suggests that the leptin receptor (LepR) signaling system, residing within the agouti-related protein (AgRP) neurons of the hypothalamus, critically contributes to obesogenic feeding, nutrient partitioning, and energy metabolism. However, the neural circuit mechanism underlying the leptin-dependent control of obesogenic feeding and energy balance remains largely elusive. Here, we show that two distinct subgroups of LepR-expressing AgRP neurons send non-collateral, GABAergic projections to the dorsomedial hypothalamic nucleus (DMH) and to the medial part of the medial preoptic nucleus (MPO) for the differential control of metabolic homeostasis and obesogenic feeding, respectively. We found that the AgRP^LepR^-DMH neural circuit plays a significant role in leptin-dependent control of metabolic homeostasis through the α3-containing GABA_A_ receptor signaling on the melanocortin 4 receptor neurons within the DMH (MC4R^DMH^). In contrast, the AgRP^LepR^-MPO neural circuit elicits dominant effects on the appetitive response to high-fat diet through the α2-containing GABA_A_ receptors on the MC4R^MPO^ neurons. Consistent with these behavioral results, the post-synaptic GABA_A_ neurons located within the DMH and MPO displayed differentiated firing responses under various feeding and nutrient conditions. Our results demonstrate that these novel GABAergic neural circuits exert differentiated control of metabolic hemostasis and obesogenic feeding via distinct post-synaptic targets of leptin-responsive AgRP neurons. The findings of two genetically and anatomically distinct GABA_A_ receptor signaling pathways within the DMH and MPO would undoubtedly accelerate the development of targeted, individualized, anti-obesity therapy.

## INTRODUCTION

Diet-induced obesity (DIO) drastically increases susceptibility to the development of metabolic, cardiovascular, and neurological diseases, as well as the recent coronavirus disease ^1–5^. Leptin, a hormone secreted by white adipocytes, interacts with the brain to communicate peripheral fuel status, suppress appetite following a meal, promote energy expenditure, and maintain blood glucose stability^6–12^. One of the key pathological characteristics in DIO is the impaired metabolic homeostasis^13,14^. There is much evidence the central leptin signaling plays a vital role in the regulation of body weight, primarily via action upon the arcuate nucleus (ARC)^7,8,15–20^. It has been found that the long-form leptin receptors are abundantly expressed in the AgRP neurons of the ARC^18,21–24^. Previous studies demonstrate that, under the physiological condition, leptin directly inhibits the LepR-expressing AgRP neurons^25,26^. However, these dynamic effects are blunted in obesity mice occurring within primary targeting neurons of the leptin signaling ^27^. On the other hand, the neural circuit and transmitter signaling mechanism underlying leptin-mediated body weight and energy balance have not yet been explored.

AgRP neurons play fundamental roles in regulating feeding behavior and body weight by releasing inhibitory NPY, AgRP and GABA transmitters into broad downstream brain areas^28–30^. The neural circuits comprised of AgRP neurons, the neural and hormonal signaling afferent to AgRP neurons, and their postsynaptic targets have been identified as key players in the regulation of energy balance and systemic insulin sensitivity^31–36^. Leptin acts to decrease food intake and promote energy expenditure by suppressing the activity of AgRP neurons^25,26,37,38^. Selective ablation of LepR in AgRP neurons gives rise to an obese phenotype and diabetes ^18,39^. Although the LepR signaling within AgRP neurons has been implicated as the dominant component for the regulation of metabolic homeostasis, the underlying neural circuit mechanism is so far poorly understood. We suggest that identification of the critical transmitter signaling components underlying the leptin-responsive neural circuit is crucial for the development of more efficient therapeutics for obesity.

Emerging data suggest a critical role of the GABA signaling on the control of feeding behavior and energy homeostasis ^20,40–55^. Numerous studies suggest that central GABA_A_ and GABAB receptor signaling exert prominent influences on feeding behavior under various metabolic states ^56–62^. For example, pharmacological activation of GABA_A_ receptor signaling in the hindbrain parabrachial nucleus (PBN) enhances the positive hedonic perception of tastes and foods, thereby promoting food intake and motivational response to food reward ^56,57,60,63–66^. We have previously shown that loss of GABA from AgRP neurons resulted in glucose intolerance^51^. Our recent work shows that the α5-containing GABA_A_ receptor signaling within the bed nucleus of the stria terminalis neurons reciprocally regulates mental disorders and obesity, implicating that the GABA_A_ receptor signaling exerts a role in controlling obesity and comorbid diseases ^30^.

In this report, we examined the neural circuit mechanism underlying the leptin action upon the AgRP neurons using a newly established and robust method by rapid inactivation of the LepR signaling within AgRP neurons^51^. We found that the LepR in the AgRP neurons plays a pivotal role in the control of obesity and metabolic homeostasis. Moreover, a subset of leptin-responsive AgRP neurons sent GABAergic projections to a group of α3-GABA_A_-expressing neurons in the DMH for regulation of leptin-mediated metabolic homeostasis. Another subset of LepR-expressing AgRP neurons innervated a group of α2-GABA_A_-expressing neurons in the MPO, which mediates leptin’s control on obesogenic feeding. Taken together, these findings suggest that the identification of key GABA_A_ signaling pathways within two distinct post-synaptic targets of a novel leptin-responsive GABAergic neural circuit will fundamentally accelerate the development of efficient treatments for obesity.

## RESULTS

To test the role of leptin receptor (LepR) in AgRP neurons, we applied our newly developed conditional knockout approach by generating two different lines of mice: *Agrp*^*nsCre*/+^::*Lepr*^*lox/lox*^::*Jax2356*^*NeoR*/+^::*Rosa26*^*tdTomato*^ mice, termed the knockout group (*Agrp::Lepr*^*KO*^), and *Agrp*^+/+^::Lepr^*lox/lox*^::*Jax2356*^*NeoR*/+^::*Rosa26*^*tdTomato*^ mice, termed the control group^51^. Immunohistochemistry and qPCR results showed the expression of tdTomato and deletion of *Lepr* in AgRP neurons 4 days after treatment of NB124, a synthetic nonsense-suppressor that can rapidly restore the functions of Cre recombinase within the AgRP neurons (Fig.1a-f). *Agrp::Lepr*^*KO*^ mice showed a significant increase in feeding and body weight one week after deletion of LepR signaling from AgRP neurons (Fig.1g,h). Furthermore, along with the increase of body weight and food intake, *Agrp::Lepr*^*KO*^ mice exhibited impaired glucose tolerance (Fig.1i). Intriguingly, chronic infusion of bicuculline (4 ng, a GABA_A_ receptor antagonist) into the 3^rd^ ventricle obviously abolished the hyperphagia responses in *Agrp::Lepr*^*KO*^ mice (Fig.1j), suggesting that facilitating the post-synaptic GABA_A_ signaling to the AgRP neural circuit is important in regulating leptin-mediated overfeeding. Meanwhile, we calculated the feeding efficiency which is presented as mg of body weight gain/kcal consumed in the mice. The feeding efficiency was significantly higher in *Agrp::Lepr*^*KO*^ mice but can be rescued by infusion of bicuculline (Fig.1k). We further showed that the expansion of white adipose tissue (WAT) accounted for the entire body weight gain in *Agrp::Lepr*^*KO*^ mice (Fig.1l). Respiratory quotient (RQ) is defined as the volume of CO_2_ released over the volume of O_2_ absorbed during respiration (defined as ratio of V_CO2_/V_O2_), which provides an indication of the nature of the substrate being used by an organism (i.e., RQ=1 for glucose utilization RQ=0.7 for lipid utilization). We observed a significant decrease in RQ from the *Agrp::Lepr*^*KO*^ mice that can be further normalized by potentiating the GABA_A_ signaling (Fig.1m). In line with other studies, these results suggest that GABA is a crucial signaling molecule by which AgRP neurons control adiposity and nutrient utilization ^47,67,68^. Further, the control and *Agrp::Lepr*^*KO*^ mice were analyzed for their glucose tolerance. The results showed that ablation of LepR in the AgRP neurons impaired glucose tolerance, which can be fully restored by infusion of GABA_A_ antagonist (Fig.1n). These results indicate that leptin regulates appetite and energy metabolism via the post-synaptic GABA_A_ signaling within the AgRP neural circuit.

**Fig. 1:**
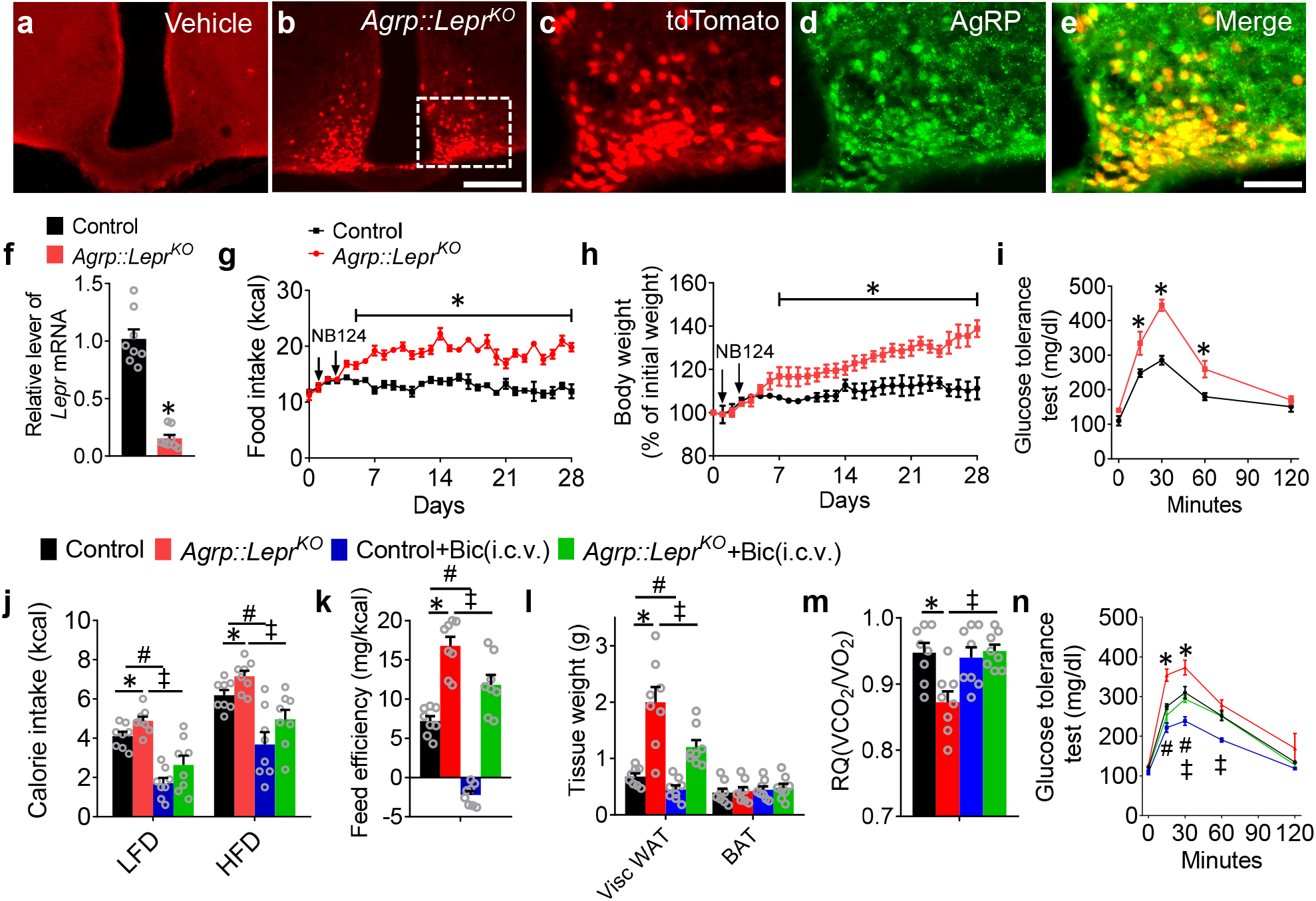
Enhancing the GABA_A_ signaling reverses obesogenic feeding and metabolic dysfunction induced by leptin deficiency in AgRP neurons. **a**,**b**, Expression of tdTomato in the arcuate nucleus after i.c.v. injection of vehicle (control mice, **a**) or NB124 (*Agrp::Lepr*^*KO*^ mice, **b**) into the 3^rd^ ventricle of *Agrp*^*nsCre*^::*Lepr*^*lox/lox*^::*Rosa26*^*tdTomato*/+^::*Jax2356*^*Neo*^ mice. Scale bar in **b** for **a** and **b**, 200 μm. **c-e**, Colocalization of tdTomato (**c**) and AgRP (**d**) in ARC. Scale bar in **e** for **c-e**, 200 μm. **f**, Real-time qPCR analysis of *Lepr* transcript levels within the AgRP neurons of the control and *Agrp::Lepr*^*KO*^ mice. (n = 8 per group; *p < 0.05). **g-I**, Daily calorie intake (**g**), body weight (**h**), and GTT (**i**) of the control and *Agrp::Lepr*^*KO*^ mice. (n = 8 per group; *p < 0.05). **j**, The 4-hr food intake by the control and *Agrp::Lepr*^*KO*^ mice after chronic infusion of bicuculline into the 3^rd^ ventricle for 4 weeks. **k**, Feeding efficiency is presented as mg of body weight gain/kcal consumed in the mice as described in **j. l**, Weight of adipose tissues in the mice as described in **j** (Visc WAT: visceral white adipose tissue; BAT: brown adipose tissue). **m**, Average value for RQ (defined as ratio of V_CO2_/V_O2_) tested on Day 28 in the mice as described in **j**. **n**, The GTT was performed in the mice as described in **j**. (n = 8 per group in **j-n**; *p < 0.05, ^#^p < 0.05, ^‡^p < 0.05). Error bars represent mean ± SEM. unpaired two-tailed t test in **f**; one-way ANOVA and followed by Tukey comparisons test in **j-l**; two-way ANOVA and followed by Bonferroni comparisons test in **g-i**, and **n**.

As in our initial step to identify the key downstream targets of AgRP neurons that contribute to the leptin-mediated responses, we examined the *Npas4* expression profile in those neurons projected by AgRP neurons. Traditional immediate early genes are regulated by neuromodulators via cAMP, neurotrophins and other paracrine factors, and their kinetics are relatively slow. In contrast, *Npas4* has a more dynamic response to activity-dependent signaling via Ca^2+^, which expressed in an activity-dependent manner not only in excitatory neurons but also in all inhibitory neurons^69^. The qPCR results showed that rapid deletion of LepR signaling from AgRP neurons resulted in a significant decrease of neural activities within the DMH and MPO (Fig.2a). Next, we devised an assay to further identify the functional relevance of each downstream target to leptin signaling. To achieve this goal, we administered DT into neonatal *Agrp*^*DTR*^ mice to ablate all AgRP neurons and examined the neural activities of downstream targets of AgRP neurons in response to leptin (Fig.2b)^70,71^. Without ablation of AgRP neurons, we found that leptin induced robust Fos induction in almost all major targets of AgRP neurons^47,48,72^. However, neonatal ablation of AgRP neurons significantly diminished the leptin-mediated Fos induction within post-synaptic neurons residing in the MPO and DMH (Fig.2c-f). These results indicate that the DMH and MPO could be the key downstream targets that critically contribute to the regulation of leptin-mediated appetitive and metabolic responses.

**Fig. 2:**
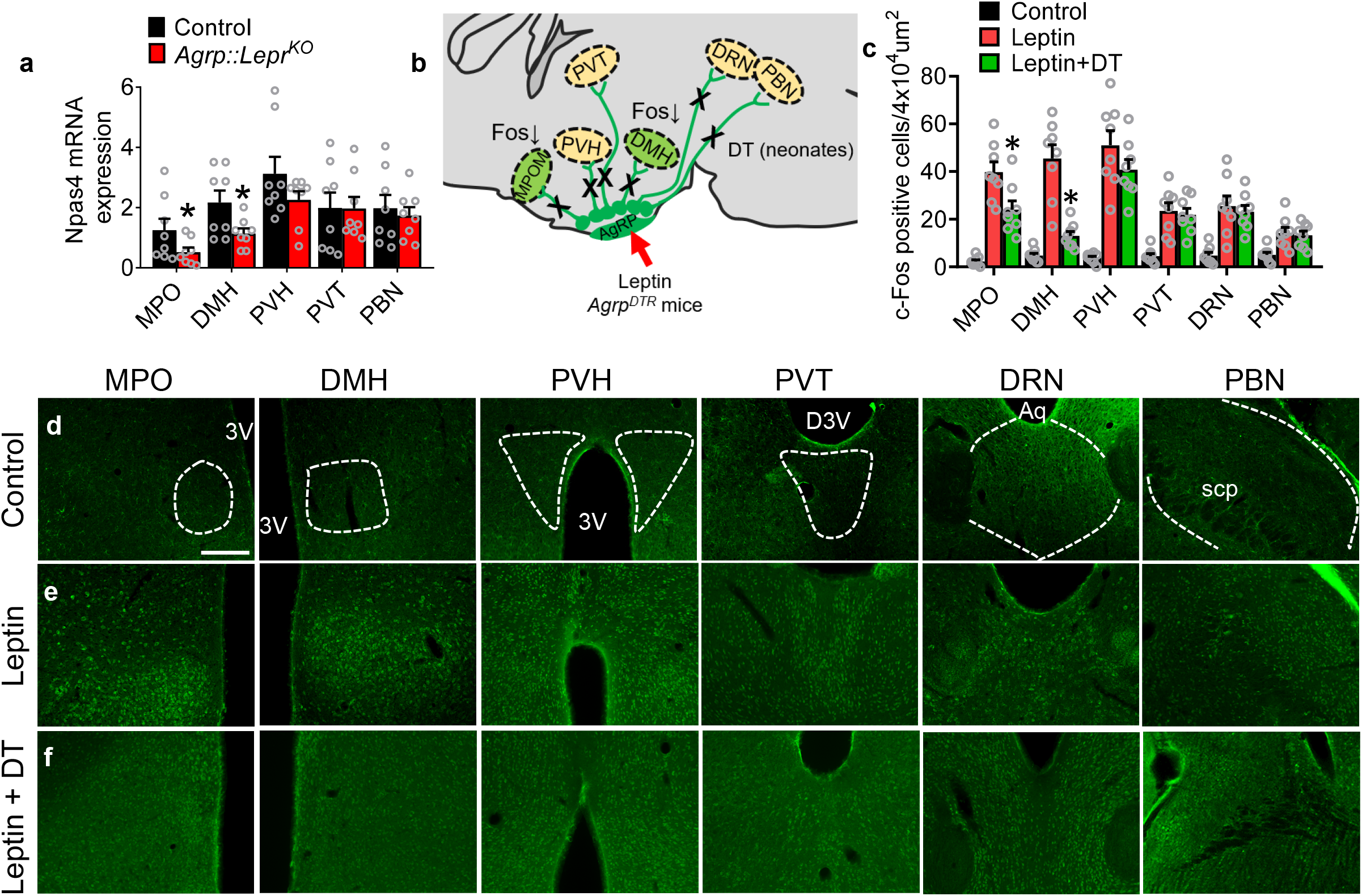
Identification of downstream targets of the leptin-responsive AgRP neurons. **a**, Real-time qPCR analysis of transcript levels of *Npas4* expressed in the MPO, DMH, PVH, PVT, and PBN in the control and *Agrp::Lepr*^*KO*^ mice. (n = 8 per group; *p < 0.05). **b**, The schematic diagram showing Fos activities in downstream targets of AgRP neurons after treatment of leptin combined with or without neonatal ablation of AgRP neurons. **c-f**, Expression profiling of Fos (green) in the MPO, DMH, PVH, PVT, DRN, and PBN under vehicle (**d**), leptin (i.p.) (**e**), leptin (i.p.) and DT (s.c. neonates) (**f**). Scale bar in **d** for **d-f**, 200 μm. (n = 8 per group; *p < 0.05). Error bars represent mean ± SEM; unpaired two-tailed t test in **a**; one-way ANOVA and followed by Tukey comparisons test in **c**.

To establish a role of the DMH and MPO within the leptin-responsive AgRP neural circuit, we applied a transsynaptic tracer to characterize the neurons in the DMH and MPO that are innervated by AgRP neurons (Fig.3a,b)^29,30,73,74^. The results derived from RNA in situ hybridization combined with transsynaptic tracing suggest that AgRP neurons make extensive connections with MC4R neurons in the DMH and MPO (Fig.3a-d and Extended Data Fig.1). Next, we performed whole-cell, patch-clamp recordings to investigate the synaptic connectivity between channelrhodopsin-2 (ChR2)-expressing AgRP axons and ZsGreen-labeled postsynaptic neurons within the DMH (Fig.3a). To test whether this connectivity was monosynaptic input from AgRP axon terminals, we perfused tetrodotoxin (TTX) and 4-aminopyridine (4-AP) into the bath to remove any network activity. We observed that inhibitory postsynaptic currents (IPSCs) in the DMH neurons triggered by photostimulation of ChR2-expressing axonal terminals of AgRP neurons were fully blocked by bicuculline (Fig.3e,f), confirming that these terminals were releasing GABA. In the presence of DNQX (a competitive AMPA/kainate receptor antagonist), AP5 (a selective NMDA receptor antagonist) and bicuculline, photostimulation of the ChR2-expressing AgRP terminals resulted in robust inhibition of action potential in postsynaptic ZsGreen neurons of the DMH in a reversible manner with significant reduction of firing rate and resting membrane potential (Fig.3g-j). We further examined the effect of leptin on the firing of postsynaptic ZsGreen neurons in the DMH. We found that leptin treatment significantly enhanced neural activities of DMH neurons (Fig.3k,l). To reveal the hierarchical structure of this neural circuit, results derived from HSV-based dual retrograde tracing showed that two distinct subgroups of AgRP neurons project to the DMH and MPO, respectively (Fig.3m-p). Taken together, these results implicated potential functional segregation of the bifurcating AgRP→DMH/MPO neural circuit.

**Fig. 3:**
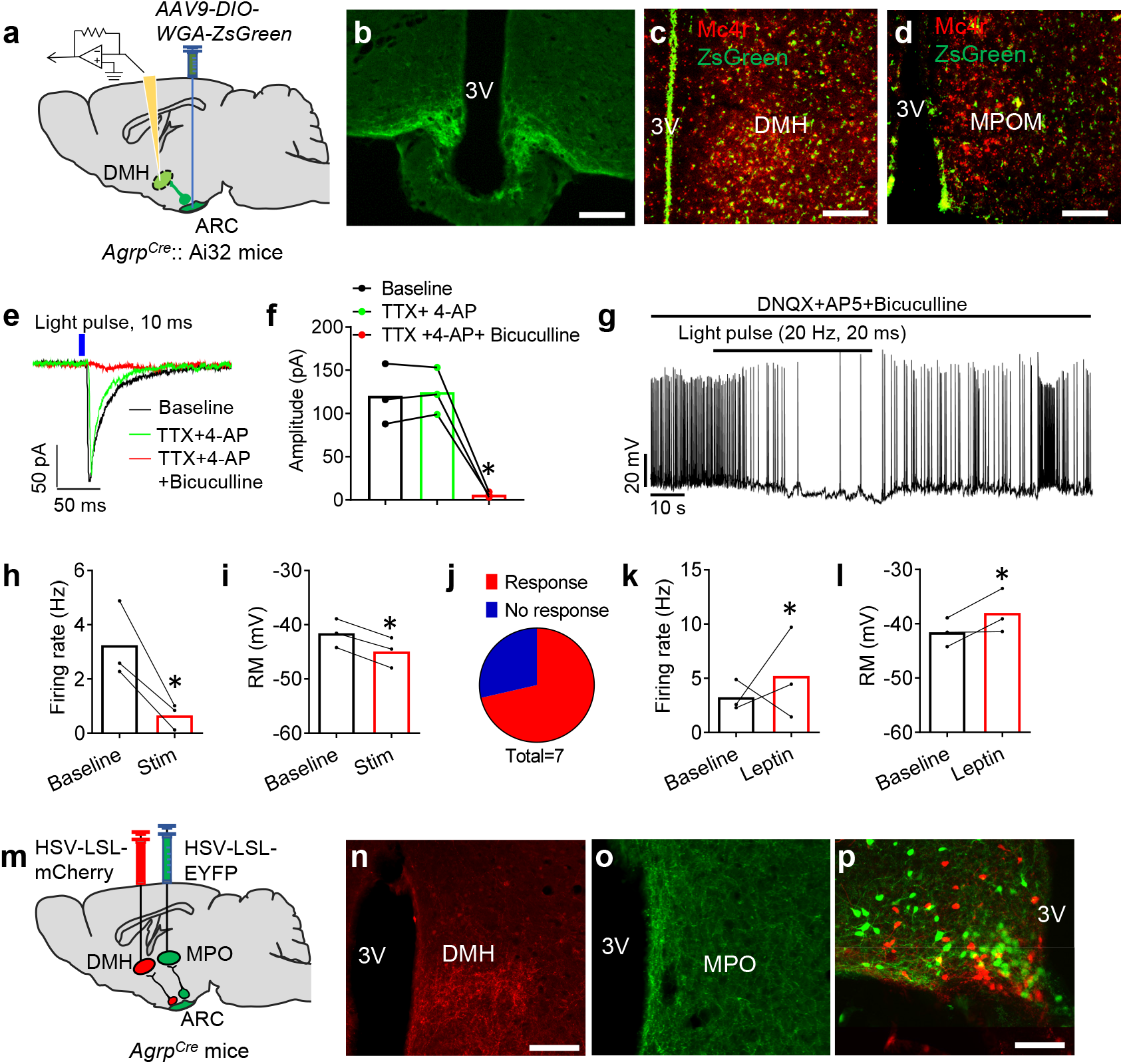
Leptin-responsive AgRP neurons directly innervate GABAergic neurons within the DMH and MPO. **a**, Diagram of transsynaptic tracing downstream targets from AgRP neurons. *AAV9-DIO-WGA-ZsGreen*, a Cre-dependent tracer that labels neurons with synaptic connections, was injected into the ARC of *Agrp*^*Cre*^::Ai32 mice. Ai32 mice express ChR2/EYFP fusion protein conditioned Cre-mediated recombination. **b**, ChR2-EYFP-labeled AgRP neurons in the ARC. Scale bar, 200 μm. **c,d**, Double-labeling of MC4R neurons with transsynaptic WGA-ZsGreen in the DMH (**c**, green; WGA-ZsGreen; red, *Mc4r* RNA in situ) and MPO (**d**, green; WGA-ZsGreen; red, *Mc4r* RNA in situ). Scale bars in **c** and **d**, 200 μm. **e**, The response of IPSCs recorded from an ZsGreen+ neuron upon photostimulation of AgRP terminals within the DMH (10 ms pulse) with a pretreatment of vehicle or TTX+4-AP, and in slices pretreated with either TTX+4-AP or TTX+4-AP+bicuculline. **f**, Statistical amplitude analysis of IPSCs from neurons recorded in **e**. (n = 3 neurons; *P < 0.05 between TTX+4-AP and TTX+4-AP+bicuculline). **g**, Representative spikes of ZsGreen-labeled DMH neurons before and after blue light pulses (20 ms/pulse, 20 Hz) shined onto AgRP axonal fibers. **h-j**, Firing frequency (**h**), resting membrane (**i**), and statistical analysis (**j**) of neurons recorded in **g**. (n < 3 neurons; *P < 0.05). **k,l**, Firing frequency (**k**), and resting membrane (**l**) of ZsGreen-labeled DMH neurons with or without leptin i.p. treatment. (n < 3 neurons; *P < 0.05). **m**, Diagram shows dual retrograde tracing of AgRP^ARC→DMH^ neurons and AgRP^ARC→MPO^ neurons by simultaneous injection of *HSV-LSL-mCherry* into the DMH and *HSV-LSL-EYFP* into the MPO of *Agrp*^*Cre*^ mice. **n-p**, The fluorescence in the DMH (**n**) and MPO (**o**), and the ARC (**p**; red, DMH-projecting AgRP neurons; green, MPO-projecting AgRP neurons). Scale bar in **n** for **n** and **o**, 200 μm; scale bar in **p**, 100 μm. Error bars represent mean ± SEM. paired two-tailed t test in **h, i, k**, and **l**; one-way ANOVA and followed by Tukey comparisons test in **f**.

To examine the role of the AgRP → DMH circuit in regulation of feeding and metabolism, we performed optogenetic manipulation in *Agrp*^*Cre*^::Ai32 mice where ChR2-eYFP was selectively expressed in AgRP neurons and axonal terminals (Fig.4a)^29,30^. Photostimulation of AgRP fibers in the DMH promoted feeding of chow diet but not high-fat diet (HFD), coupled with disrupted glucose intolerance (Fig.4b,c). Consistently, photostimulation of post-synaptic MC4R^DMH^ neurons resulted in a significant decrease in feeding of chow but not HFD and improved glucose intolerance (Fig.4d,e). Bilateral infusion of bicuculline into the DMH of *Agrp::Lepr*^*KO*^ mice resulted in reduced chow feeding and a significant reduction of body weight, coupled with improved glucose tolerance (Fig.4f-i). Moreover, our studies showed that blockage of GABA_A_ receptor in the DMH significantly rescued the feeding efficiency, expansion of WAT, and the RQ profiles in *Agrp::Lepr*^*KO*^ mice (Fig.4j-l). These results suggest that GABA_A_ receptor signaling within the AgRP→DMH circuit is critical for the regulation of chow diet intake, adiposity, and nutrient utilization.

**Fig. 4:**
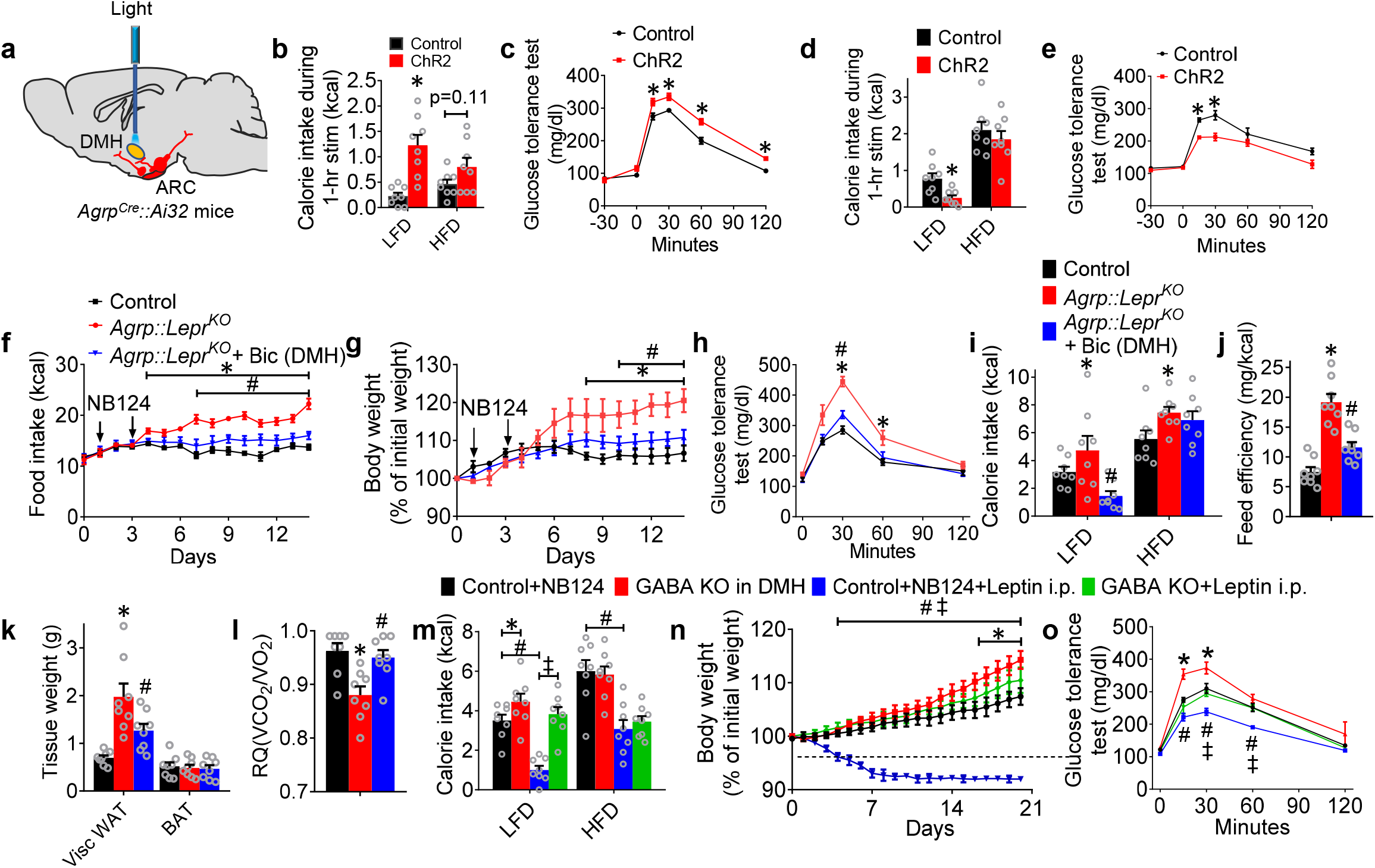
The leptin-responsive AgRP-DMH circuit regulates glucose tolerance, and nutrient partitioning. **a**, Diagram showing the photostimulation of AgRP axonal terminals within the DMH of *Agrp*^*Cre*^::Ai32 mice. **b,c**, 1-hr food intake of LFD and HFD (**b**) and GTT (**c**) after photostimulation of the AgRP→DMH circuit. (n = 8 per group; *p < 0.05; p = 0.11 between Control and ChR2 in HFD group). **d,e**, 1-hr food intake of LFD and HFD (**d**) and GTT (**e**) after photostimulation of the MC4R^DMH^ neurons. (n = 8 per group; *p < 0.05). **f-h**, Daily food intake (**f**), body weight (**g**) and GTT (**h**) in control mice, *Agrp::Lepr*^*KO*^ mice with bicuculline or vehicle injected into the DMH. The saline or bicuculline (0.1 ul/min, 5min) was chronically infused into the DMH via the osmotic minipump. (n = 8 per group; *p < 0.05, ^#^p < 0.05). **I**, 4-hrs food intake of LFD and HFD (**f**) in the mice described in **f-h**. **j**, Feed efficiency in the mice described in **f-h. k**, Weight of adipose tissues in the mice described in **f-h. l**, Average value for RQ (V_CO2_/V_O2_) in the mice described in **f-h**. (n = 8 per group in **i-l**; *p < 0.05, ^#^p < 0.05). **m-o**, 4-hrs food intake of LFD and HFD (**m**), body weight (**n**) and GTT (**o**) in control mice, GABA KO mice with or without treatment of leptin (1 μg/g body weight). The *AAV9*-*fDIO-WGA-nsCre-mCitrine* virus or control *AAV9-fDIO-WGA-mCitrine* virus was injected into the ARC of *Npy*^*Flp*^::*Gad1*^*lox/lox*^::*Gad2*^*lox/lox*^::*Rosa26*^*tdTomato*^ mice followed with injection of NB124 into the DMH. (n = 8 per group in **m-o**; *p < 0.05, ^#^p < 0.05, ^‡^p < 0.05). Error bars represent mean ± SEM. unpaired two-tailed t test in **b** and **d**; one-way ANOVA and followed by Tukey comparisons test in **b, d, f, k**; two-way ANOVA and followed by Bonferroni comparisons test in **c-h, n**, and **o**.

To better understand the contribution of the AgRP→DMH circuit to the control of leptin-mediated functions, we bilaterally injected the *AAV9-fDIO-WGA-nsCre* virus into the ARC of *Npy*^*Flp*^::*Gad1*^*lox/lox*^::*Gad2*^*lox/lox*^::*Rosa26*^*tdTomato*^ mice followed with treatment of NB124 into the DMH, a strategy which can specifically inactivate the GABA signaling from those DMH neurons directly innervated by AgRP neurons. We showed that acute ablation of GABA signaling from the AgRP-projected DMH neurons significantly increased feeding of chow diet without affecting intake of HFD (Fig.4m). Under the treatment of chow diet, these GABA signaling-deficient mice displayed a moderate but significant increase in body weight coupled with impaired glucose intolerance (Fig.4n,o). More importantly, inactivation of post-synaptic GABA signaling in the AgRP → DMH circuit blunted the actions of systemically administered leptin on chow diet feeding, body weight, and glucose tolerance (Fig.4m-o). Overall, these data demonstrate that the GABA signaling within the downstream targets of the AgRP→DMH neural circuit plays a significant role in control of leptin-mediated metabolic homeostasis.

To better understand the physiological responses of these GABAergic neurons in the context of feeding and metabolic regulation, we employed *in vivo* opto-tetrode system to reveal the dynamic activities of MC4R^DMH^ neurons^73^. With the injection of *AAV2-DIO-ChR2-GFP* into the DMH of *Mc4r*^*Cre*^ mice, the ChR2 neurons can be identified by the latencies of evoked spikes accurately following high-frequency photostimulation, as well as the identical waveforms of evoked and spontaneous spikes^73^. We investigated the activities of MC4R^DMH^ neurons and non-MC4R^DMH^ neurons under low-fat diet (LFD), HFD, and hyperglycemia condition. A total 13 MC4R^DMH^ neurons and 9 non-MC4R^DMH^ neurons were identified through optogenetic-invoked spikes. The results showed that 7 out of 13 identified MC4R neurons responded to the feeding with LFD, with a reduction of firing rate from 22.4 Hz to 12.7 Hz (Fig.5a,c). We also observed 6 out of 13 MC4R neurons that respond to the enhancement of blood glucose with a reduction of firing rate from 21.9 Hz to 13.9 Hz (Fig.5b,d). This result indicates that the MC4R^DMH^ neurons mediate feeding and glucose tolerance.

**Fig. 5:**
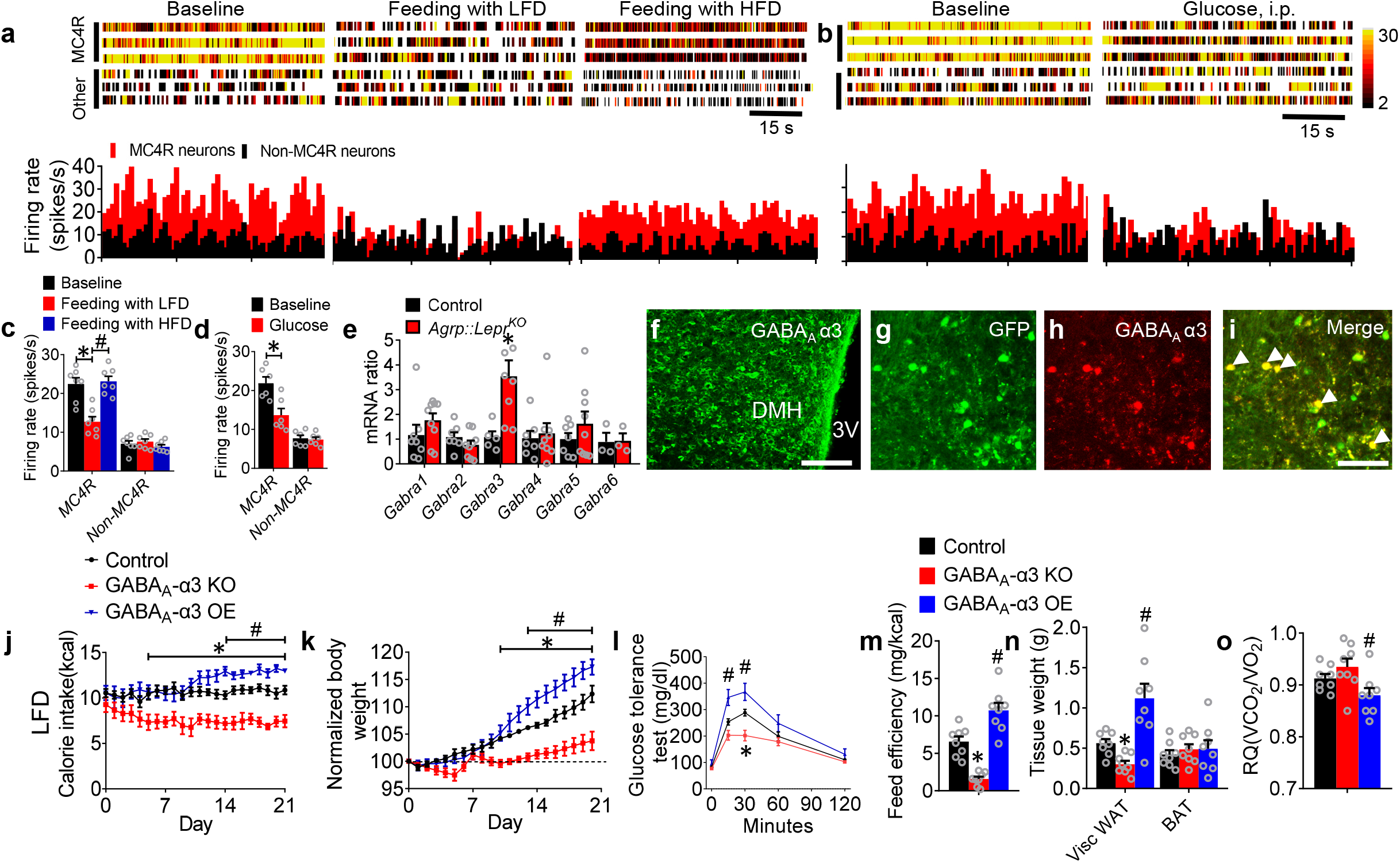
The α3-GABA^A^ receptor signaling within the MC4R^DMH^ mediates glucose tolerance and nutrient partitioning. **a-d**, Raster plots and firing rate of representative MC4R^DMH^ neurons and non-MC4R^DMH^ neurons when mice under baseline, feeding with LFD and HFD (**a**), and under glucose i.p. injection (**b**). The statistical analysis of the firing rate under baseline, feeding with LFD and HFD (**c**), and under glucose treatment (**d**) were calculated. (n = 6 or 7 in each group; *p < 0.05, ^#^p < 0.05). **e**, Expression level of *Gabra1-Gabra6* in the DMH neurons of control and *Agrp::Lepr*^*KO*^ mice. **f**, Immunostaining showing the expression of GABA_A_ α3 in the DMH. Scale bar, 200 μm. **g-I**, Colocolization of GABA_A_ α3 and transsynaptically labeled neurons by WGA-ZsGreen in the DMH after injection of *AAV9-DIO-WGA-ZsGreen* into the ARC of *Agrp*^*Cre*^ mice. **j-l**, The intake of LFD (**j**), body weight (**k**) and GTT (**l**) were performed in *Mc4r*^*Cre*^::*Rosa26*^*Cas9*^ mice with a bilateral injection of vehicle, *AAV9-DIO-Gabra3*^*sgRNA*^-*tdTomato* (GABA_A_-α3 KO), or *AAV9-DIO-Gabra3*^*cDNA*^-*tdTomato* (GABA_A_-α3 OE) into the DMH. **m**, Feeding efficiency in the mice described in **j-l. n**, Weight of adipose tissues in the mice described in **j-l. o**, Average value for RQ (V_CO2_ /V_O2_) in the mice described in **j-l**. (n = 8 per group in **j-o**; *p < 0.05, ^#^p < 0.05). Error bars represent mean ± SEM. unpaired two-tailed t test in **d** and **e**; one-way ANOVA and followed by Tukey comparisons test in **c, m**, and **n**; two-way ANOVA and followed by Bonferroni comparisons test in **j-l**.

Our results showed that GABA_A_ receptors in the DMH are involved in the regulation of feeding and nutrient partitioning; thus, we attempted to identify the key GABA_A_ receptor subunits that are functionally relevant to glucose and feeding regulation. Results of qPCR analysis showed that, among all major regulatory α subunits, the transcript level of *Gabra3* (encoding GABA_A_ receptor α3 subunits) in the DMH neurons of *Agrp::Lepr*^*KO*^ mice was robustly enhanced (Fig.5e). Immunostaining data showed the α3-GABA_A_ signaling was abundantly expressed within the DMH (Fig.5f). Our transsynaptic tracing study further confirmed that the α3-GABA_A_ signaling was highly co-localized within the post-synaptic targets of the AgRP→DMH neural circuit (Fig.5g-i). To understand the functional roles of the α3-GABA_A_ signaling with the downstream DMH neurons in relevancy to leptin-mediated feeding and metabolism, *Mc4r*^*Cre*^::*Rosa26*^*Cas9*^ mice were injected with *AAV9-Gabra3*^*sgRNA*^-*mCherry* into DMH. We found that knockout of *Gabra3* signaling in the MC4R^DMH^ neurons reduces feeding of chow diet and body weight while the glucose tolerance was significantly improved (Fig.5j-l and Extended Data Fig.2). Meanwhile, genetic deficiency in α3-GABA_A_ signaling significantly decreased feeding efficiency and WAT (Fig.5m-o). On the other hand, the gain-of-function study showed that overexpression of α3-GABA_A_ within the same MC4R^DMH^ neurons manifested the exact opposite phenotypes, including moderately increased chow diet feeding and body weight, significantly increased feeding efficiency and WAT adiposity, and profoundly decreased RQ (Fig.5m-o). These results suggest that the MC4R^DMH^ neurons play a significant role in control of glucose tolerance, nutrient partitioning, and feeding efficiency through the α3-GABA_A_ signaling.

Next, we characterized the functional role of the AgRP→MPO circuit in leptin-mediated feeding and metabolism. Contrasting to the role of the AgRP→DMH circuit, photostimulation of AgRP fibers within the MPO led to a significant increase in feeding of HFD without affecting the normal response to LFD or glucose intolerance (Fig.6a, and fig. S3A). Similarly, photostimulation of post-synaptic MC4R^MPO^ neurons specifically suppressed intake of HFD but not LFD while glucose tolerance remained intact (Fig. 6B and Extended Data Fig.3b). Further, by employing a similar genetic strategy described in Fig.4, M-O, we showed that acute ablation of GABA signaling from the post-synaptic MC4R^MPO^ neurons within the AgRP→MPO circuit significantly increased HFD feeding while chow diet intake and glucose tolerance were unaffected (Fig.6c and Extended Data Fig.3c). Notably, genetic disruption of post-synaptic GABA signaling within the MPO abolished the anorexic effect of leptin on HFD feeding, while the leptin-induced effect on LFD feeding remained intact (Fig.6c). Taken together, these data demonstrate that AgRP^LepR→^MPO neural circuit and post-synaptic GABAergic signaling play an exclusive role in control of leptin-mediated obesogenic feeding.

**Fig. 6:**
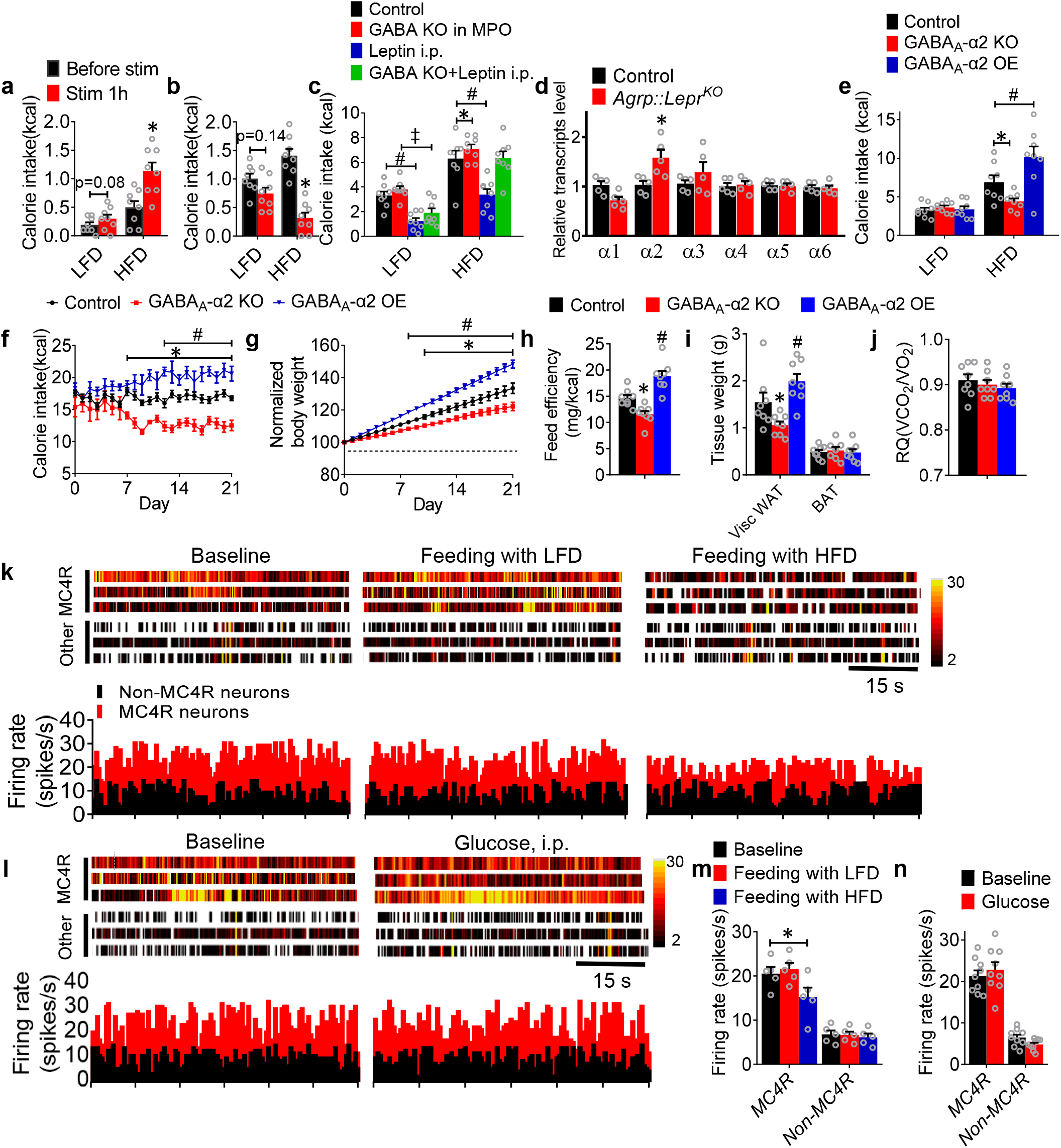
The leptin-responsive AgRP-MPO circuit regulates obesogenic feeding via the α2-GABA_A_ signaling. **a,b**, 1-hr food intake of LFD and HFD after photostimulation of the AgRP→MPO circuit (**a**) or MC4R^MPO^ neurons (**b**). (n = 8 per group; *p < 0.05). (c) 4-hrs food intake of LFD and HFD in the control, *Agrp::Lepr*^*KO*^ mice with leptin (i.p.) or vehicle. (n = 8 per group; *p < 0.05, ^#^p < 0.05, ^‡^p < 0.05). **d**, Expression level of *Gabra1 - Gabra6* in the MPO neurons of control and *Agrp::Lepr*^*KO*^ mice. **e-g**, The 4-hrs food intake of HFD (**e**), daily food intake of HFD (**f**), and body weight (**g**) were performed in *Mc4r*^*Cre*^::*Rosa26*^*Cas9*^ mice with bilateral injection of either vehicle, *AAV9-DIO-Gabra2*^*sgRNA*^-*tdTomato* (GABA_A_-α2 KO), or *AAV9-DIO-Gabra2*^*cDNA*^-tdTomato (GABA_A_-α2 OE) into the MPO. **h**, Feeding efficiency in the mice described in **e-g. i**, Weight of adipose tissues in the mice described in **e-g. j**, Average value for RQ (V_CO2_/V_O2_) of the mice described in **e-g**. (n = 8 per group in **e-j**; *p < 0.05, ^#^p < 0.05). **k-n**, Raster plots and firing rate of representative MC4R^MPO^ neurons and non-MC4R^MPO^ neurons under well-fed baseline, feeding with LFD and HFD (**k**), and under glucose i.p. injection (**l**). The statistical analysis of the firing rate under baseline, feeding with LFD and HFD (**m**), and under glucose treatment (**n**) were calculated. (n = 9 per group; *p < 0.05). Error bars represent mean ± SEM. paired two-tailed t test in **a, b**; unpaired two-tailed t test **d**; one-way ANOVA and followed by Tukey comparisons test in **c, h, i**, and **m**; two-way ANOVA and followed by Bonferroni comparisons test in **e-g**.

To identify the critical GABA_A_ receptor components contributed to these phenotypes, we examined the expression profile of the major regulatory α1-α6 subunits within the MPO. Our statistical results showed that *Gabra2* (encoding GABA_A_ receptor α2 subunits) in the MPO neurons of *Agrp::Lepr*^*KO*^ mice was significantly enhanced (Fig.6d).

To establish the physiological roles of the α2-GABA_A_ signaling in the context of leptin-mediated behavioral and metabolic phenotypes, *Mc4r*^*Cre*^::*Rosa26*^*Cas9*^ mice were injected with *AAV9-Gabra2*^*sgRNA*^-*mCherry* into the MPO. We found that knockout of *Gabra2* signaling in the MC4R^MPO^ neurons reduces feeding of HFD and body weight while the glucose tolerance was not affected (Fig.6e-g and Extended Data Fig.4). Meanwhile, genetic deficiency in α2-GABA_A_ signaling significantly decreased feeding efficiency and WAT (Fig.6h-j). On the other hand, the gain-of-function study showed that overexpression of α2-GABA_A_ within the same MC4R^MPO^ neurons manifested the exact opposite phenotypes, including moderately increased HFD feeding and body weight, significantly increased feeding efficiency and WAT adiposity (Fig.6h-j). Together, these results suggest that the α2-GABA_A_ signaling within the MC4R^MPO^ neurons exerts a key role in control of obesogenic feeding and obesity.

To better understand the functional role of MC4R^MPO^ neurons in the regulation of feeding of HFD, we employed opto-tetrode recording in the MPO to investigate the activities of MC4R^MPO^ neurons and non-MC4R^MPO^ neurons under LFD, HFD, and hyperglycemia condition. A total 9 MC4R^DMH^ neurons and 8 non-MC4R^DMH^ neurons were identified through optogenetic-invoked spikes. The results showed that 5 out of 9 identified MC4R neurons responded to the feeding with HFD, with a reduction of firing rate from 20.5 Hz to 15.2 Hz (Fig.6k,m). These 9 MC4R neurons did not respond to the enhancement of blood glucose (Fig.6l,n). In conclusion, these data indicate that the post-synaptic MC4R^MPO^ neurons within the AgRP^LepR^-MPO neural circuit mediate leptin-dependent control of hyperphagic response to the obesogenic diet.

## DISCUSSION

Leptin exerts its behavioral and metabolic effects by dynamic influences onto the neural signaling within the hypothalamic AgRP neurons that are otherwise susceptible to the obesity-induced leptin resistance. In this report, we explored the neural circuit and transmitter signaling mechanism underlying leptin-mediated feeding and energy metabolism (Fig.7). We identified and characterized a unique bifurcating GABAergic neural circuit bearing distinct functional significance: the AgRP^LepR^→DMH circuit play a critical role in control of metabolic homeostasis through the α3-GABA_A_ receptor signaling, whereas the AgRP^LepR^→MPO circuit dominantly regulates high-fat diet intake through the α2-GABA_A_ receptor signaling. These leptin-responsive neural circuits play a fundamental role in regulation of hyperphagia and metabolic dysfunction in obesity. Further, enhancement of these GABA_A_ receptor signaling systems found within the distinct post-synaptic targets exerts potent suppression of obesogenic feeding and restoration of metabolic homeostasis, a combinatorial effect which prevents obesity. We suggest that manipulation of these neural circuits and associated GABA_A_ pathways can benefit novel obesity therapeutics.

**Fig. 7:**
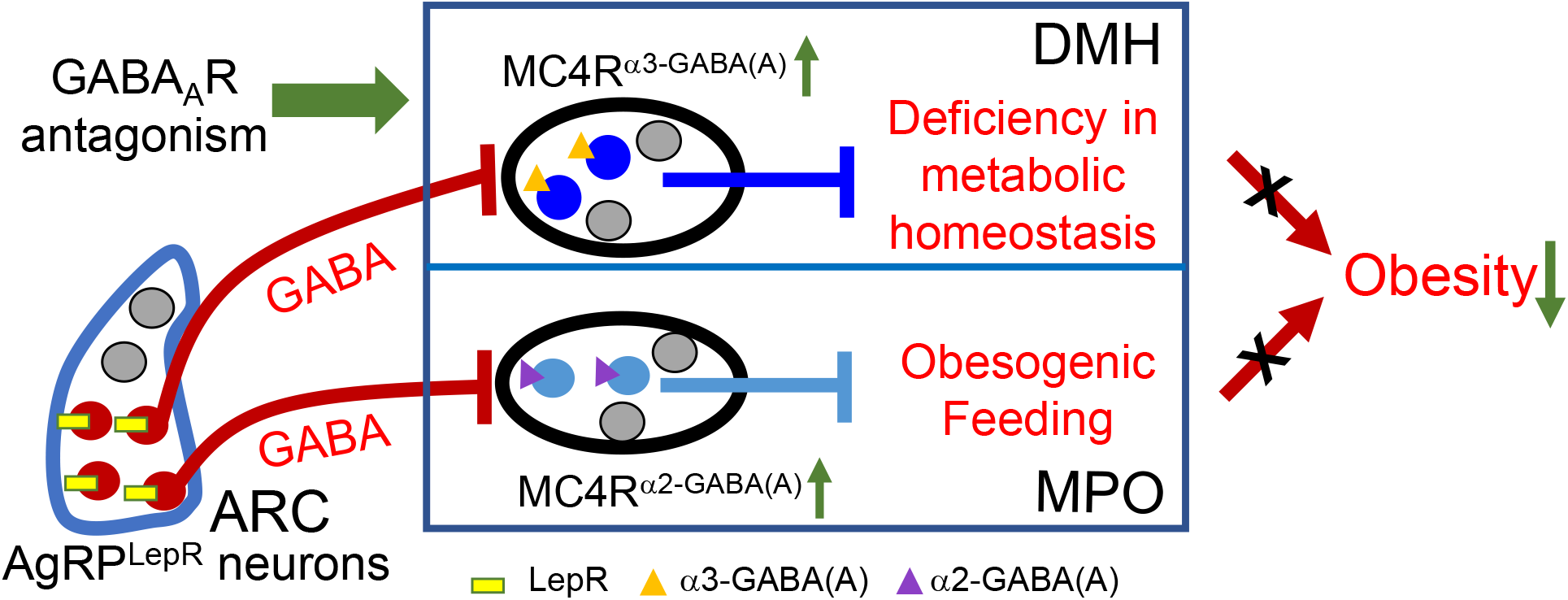
Diagram showing a GABAergic neural circuit reverses obesity by overriding central leptin resistance. The GABAergic AgRP^LepR^→DMH circuit plays a critical role in control of leptin-mediated metabolic homeostasis through the α3-GABA_A_ signaling within the MC4R^DMH^ neurons, whereas the AgRP^LepR^→MPO circuit dominantly regulates feeding of obesogenic diet through α2-GABA_A_ signaling within the MC4R^MPO^ neurons. Together, this neural circuit overrides hypothalamic leptin resistance by exerting differentiated actions onto the post-synaptic targets, a combinatorial effect which prevents obesity.

This study utilized our newly established inducible knockout strategy to achieve rapid, post-developmental, targeted inactivation of LepR signaling which precisely illuminates the pathophysiological roles of central leptin signaling on the control of nutrient partitioning, and feeding efficiency. Compared with the mild effects of the non-inducible manipulation of the brain LepR signaling, our Lepr-KO model showed a robust disruption in various feeding and metabolism parameters, culminating into severe obesity which can only be achieved by global genetic deletion or the CRISPR–Cas9 technique ^18,39,75,76^. Our technique boasts many attractive features, such as the large repertoire of conditional mouse lines, transient and non-BBB-crossing inducer, and easy combination with numerous viral tools, that can be feasible to apply for many other neurological and endocrine questions.

The profiling assay using neonatal ablation of AgRP neurons revealed key neural population responding to leptin. Among various downstream targets of the AgRP circuit, the DMH and MPO neurons are most sensitive to the AgRP-dependent leptin signaling. Acute stimulation of post-synaptic MC4R neurons within the DMH but not MPO blunted glucose tolerance and feeding with LFD not HFD. Acute stimulation of MC4R neurons within the MPO specifically blunted the feeding with HFD. Further, we developed a novel viral-mediated, circuit-dependent, gene-editing technique to specifically ablate the GABA signaling from a subpopulation of AgRP downstream targets. Our results indicated that the GABA signaling within the AgRP→DMH circuit plays a significant role in metabolic homeostasis, whereas the GABAergic signaling within the AgRP→MPO circuit specially regulate leptin-dependent obesogenic feeding.

It has been demonstrated that GABA_A_ receptor antagonists can prevent abnormalities in leptin actions on paraventricular hypothalamic neurons, which indicates GABA_A_ signaling seems to contribute to a persistently reduced negative feedback of adiposity signals in obese models. Moreover, acute bicuculline administration seems to be able to suppress food intake and prevent the obesity phenotype^77^. Further studies identified the differential roles of subunits of GABA_A_ receptors in the regulation of feeding and body weight. Studies using genetic knock-in mice revealed a strong correlation between individual GABA_A_ subunits and specific phenotypes in BDZ treatment: the α1 subunit with sedative and amnesic effects; the α2 subunit with myorelaxant effects, the α3 subunit with anxiolytic effects, and the α5 subunit with neural plasticity and metabolic effects ^57,78,79^. Another study implicated that β2/β3-containing GABA_A_ receptors contribute to feeding behavior in a hypothalamic circuit ^80^. Therefore, it is important to discover how the GABA_A_ subunit in the hypothalamus is involved in the regulation of feeding. We employed the CRISPR-Cas9 gene-editing method to comprehensively evaluate the functions of GABA_A_ subunits α3 and α2 within a leptin-responsive neural circuit. Genetic deletion of α3 in the MC4R neurons in the DMH suppressed nutrient partitioning and feeding efficiency. Genetic deletion of α2 in the MC4R neurons in the MPO inhibited obesogenic feeding. These studies demonstrate that the GABA_A_ subunits α3 and α2 play differentiated roles in the control of leptin-mediated hypometabolism and obesogenic dietary intake.

Utilizing the in vivo optrode recording system, we had the unique advantage to perform stable intracellular recordings from MC4R neurons in the DMH and MPO in an intact and awake animal, allowing us to study complex behaviors and physiological responses. Here, we focused on MC4R neurons in the DMH, which are the post-synaptic targets of AgRP neurons. Our results showed that these groups of neurons are the most relevant to hyperglycemia due to their suppressed neural activity associated with high glucose levels. This study implicated that MC4R neurons in the DMH is dynamically involved in glucose homeostasis and LFD intake. On the other hand, the MC4R neurons in the MPO could respond to HFD during feeding behavior tests. This reveals the different roles of MC4R neurons in the various downstream targets of AgRP neurons and their regulatory effects on feeding and metabolism. In conclusion, MC4R signaling located within the DMH and MPO displayed differentiated firing responses under various feeding and nutrient conditions.

In conclusion, LepR in the AgRP neurons and the associated neural circuits and receptors are the primary mediators of leptin action in obesogenic feeding and metabolic homeostasis. We suggest that novel therapeutics targeting the α2/α3-GABA_A_ receptor signaling, and associated pathways identified within distinct post-synaptic targets of the leptin-responsive GABAergic neural circuit would prevent obesity.

## ACKNOWLEDGEMENT

The Metabolomics Core and the Mouse Metabolism Core at Baylor College of Medicine provided various technical support. Some AAV vectors were packaged by the Optogenetics and Viral Design/Expression Core at Baylor College of Medicine. Cecilia Ljungberg with the RNA In Situ Hybridization Core facility at Baylor College of Medicine provided technical support on the in situ hybridization.

## Author contributions

Q.W. and Y.Han conceived and designed the experiments. Y.Han performed the surgery, *in vivo* electrophysiology, immunohistological imaging, and relevant data analysis. Y.He performed slice electrophysiology. Y.Han performed the behavioral tests and relevant data analysis. L.H. performed the colony management, genotyping. Q.W. and Y.Han wrote the manuscript with inputs from all authors on the manuscript. Y.X. provided comments for manuscript. Q.W. supervised the project.

## Data and materials availability

All data needed to evaluate the conclusions in the paper are present in the paper and/or the Supplementary Materials.

---

**Extended Data Fig. 1 Transsynaptic tracing from AgRP neurons to the MC4R neurons in the DMH and MPO. a-f**, Representative images showing the expression of WGA-ZsGreen and MC4R in the DMH of the *Agrp*^*Cre*^::Ai32 mice with *AAV9-DIO-WGA-ZsGreen* injected into the ARC. Scale bar in **a** for **a-c**, 200 μm; Scale bar in **d** for **d-f**, 100 μm. **g-l**, Representative images showing the expression of WGA-ZsGreen and MC4R in the MPO. Scale bar in **g** for **g-i**, 200 μm; Scale bar in **j** for **j-l**, 100 μm.

**Extended Data Fig. 2 GABA_A_ receptor α3 subunits in the DMH do not mediate HFD intake. a-c**, The food intake of HFD (**a**), body weight (**b**) and GTT (**c**) were performed in *Mc4r*^*Cre*^::*Rosa26*^*Cas9*^ mice with bilateral injection of either vehicle, *AAV9-DIO-Gabra3*^*sgRNA*^-*tdTomato* (a3 KO) into the DMH. (n = 8 per group; *p < 0.05). **d**, Real-time qPCR analysis of transcript levels of *a3* expressed in MC4R^DMH^ neurons isolated by fluorescence-activated cell sorting in the control and *a3* KO mice. (n = 8 per group; *p < 0.05). **e-g**, The food intake of HFD (**e**, body weight (**f**) and GTT (**g**) were performed in *Mc4r*^*Cre*^ mice with bilateral injection of either vehicle, *AAV9-DIO-Gabra3*^*cDNA*^-*tdTomato* (a3 OE) into the DMH. (n = 8 per group; *p < 0.05). **h**, Real-time qPCR analysis of transcript levels of *a3* expressed in MC4R^DMH^ neurons isolated by fluorescence-activated cell sorting in the control and *a3* OE mice. (n = 8 per group; *p < 0.05). Error bars represent mean ± SEM. one-way ANOVA and followed by Tukey comparisons test in **d** and **h**; two-way ANOVA and followed by Bonferroni comparisons test in **c** and **g**.

**Extended Data Fig. 3 AgRP→MPO neural pathway dose not regulate glucose tolerance. a,b**, GTT after photostimulation of the AgRP→MPO circuit (**a**) or MC4R^MPO^ neurons. (n = 8 per group; *p < 0.05). **c**, GTT in control mice, GABA KO mice and leptin injected mice. The saline or leptin (4 mg/kg, i.p.) was injected respectively. Food intake or GTT was tested 30 minutes later. (n = 8 per group; *p < 0.05). Error bars represent mean ± SEM.

**Extended Data Fig. 4 GABA_A_ receptor α2 subunits in the MPO do not mediate LFD intake and glucose tolerance. a-c**, The food intake of LFD (**a**), body weight (**b**) and GTT (**c**) were performed in *Mc4r*^*Cre*^::*Rosa26*^*Cas9*^ mice with bilateral injection of either vehicle, *AAV9-DIO-Gabra2*^*sgRNA*^-*tdTomato* (a2 KO) into the MPO. (n = 8 per group; *p < 0.05). **d**, Real-time qPCR analysis of transcript levels of *a2* expressed in MC4R^MPO^ neurons isolated by fluorescence-activated cell sorting in the control and *a2* KO mice. (n = 8 per group; *p < 0.05). **e-g**, The food intake of LFD (**e**), body weight (**f**) and GTT (**g**) were performed in *Mc4r*^*Cre*^ mice with bilateral injection of either vehicle, *AAV9-DIO-Gabra2*^*cDNA*^-*tdTomato* (a2 OE) into the MPO. (n = 8 per group; *p < 0.05). **h**, Real-time qPCR analysis of transcript levels of *a2* expressed in MC4R^MPO^ neurons isolated by fluorescence-activated cell sorting in the control and *a2* OE mice. (n = 8 per group; *p < 0.05). Error bars represent mean ± SEM. one-way ANOVA and followed by Tukey comparisons test in **d** and **h**.

