## Supplementary figures and images for "Reversal of obesogenic feeding and hypometabolism by a bifurcating GABAergic neural circuit"

### Supplemental Figures

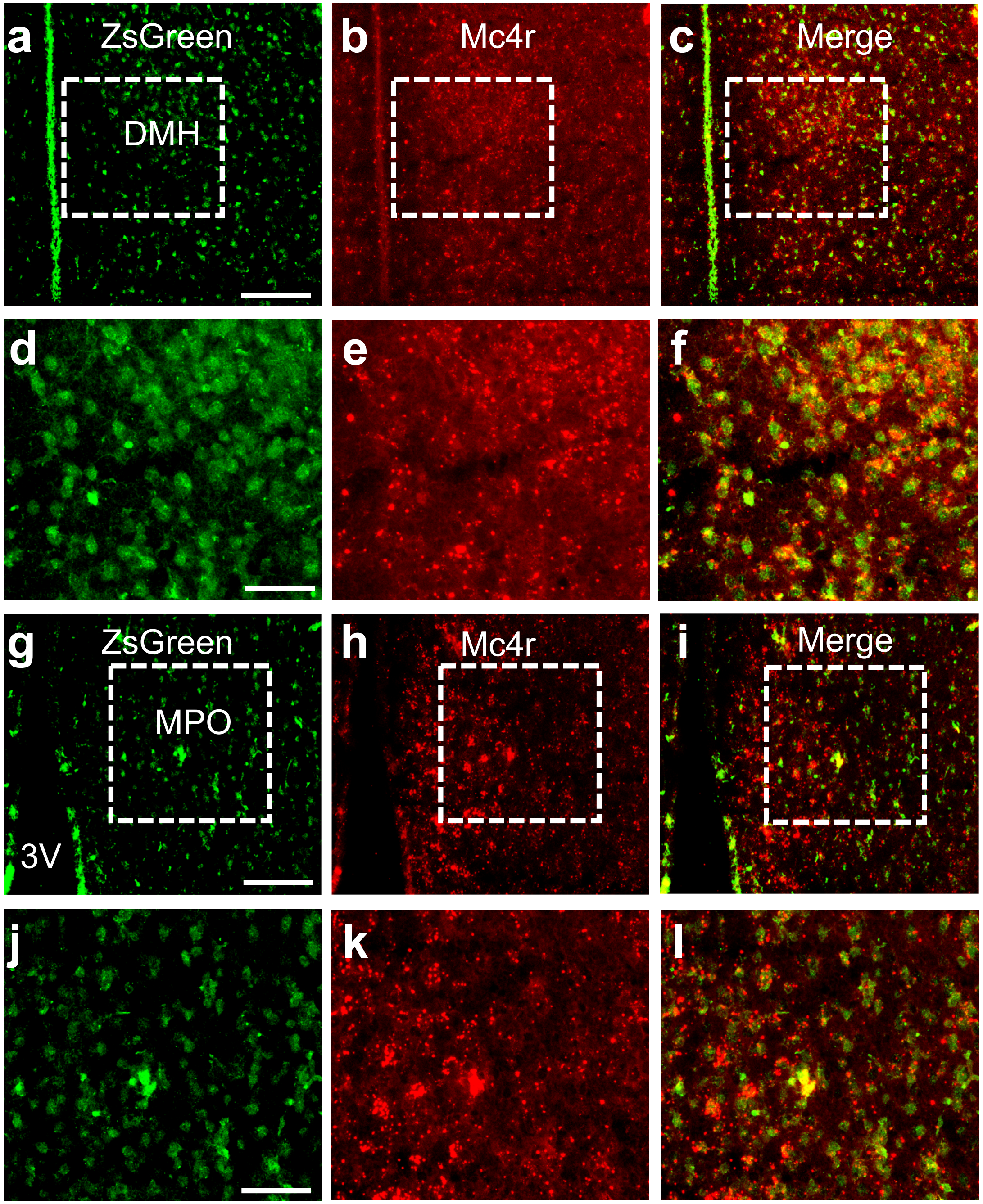


Extended Data Fig.1

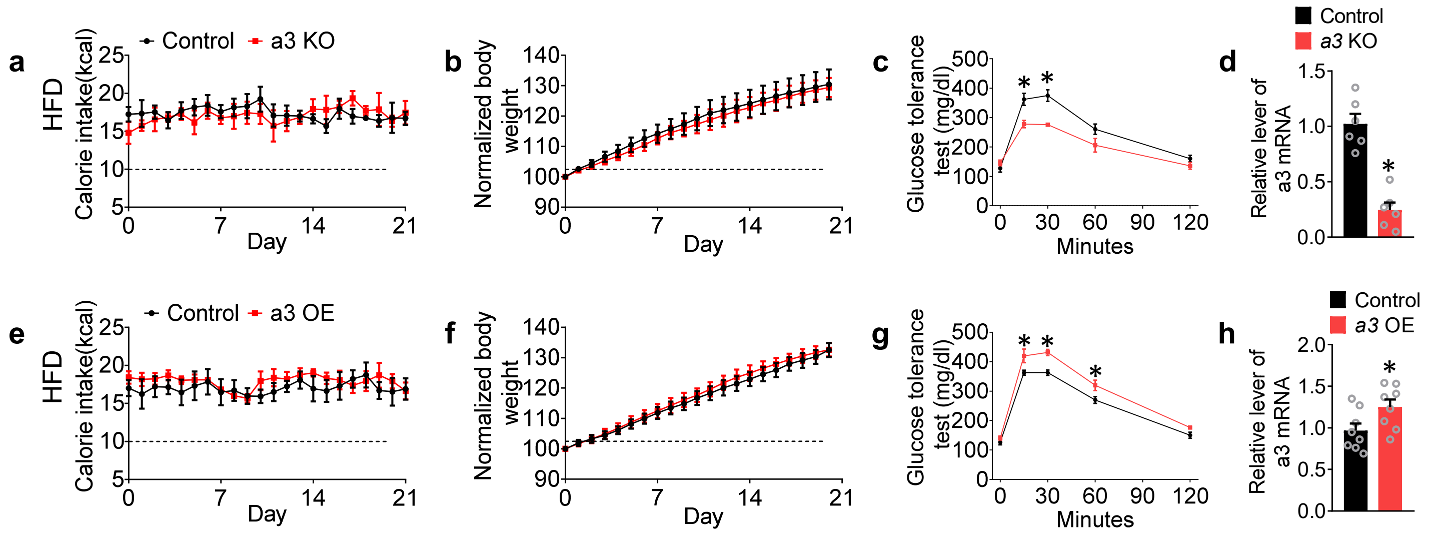


Extended Data Fig.2

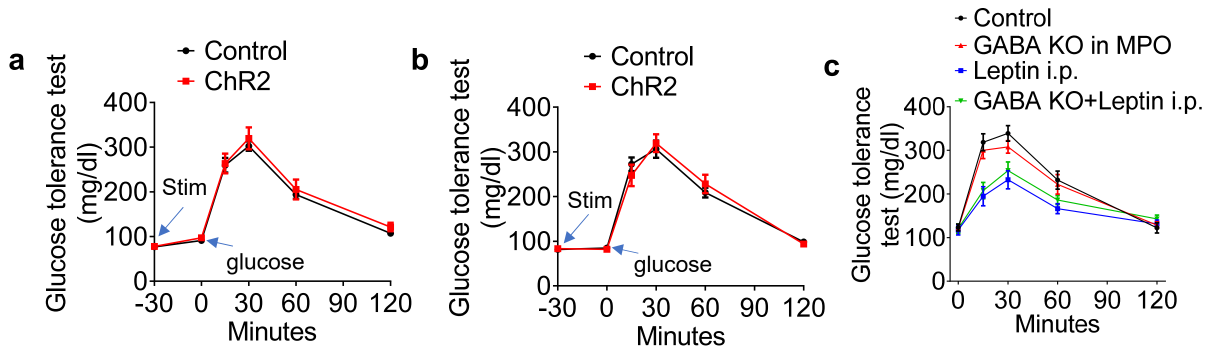


Extended Data Fig.3

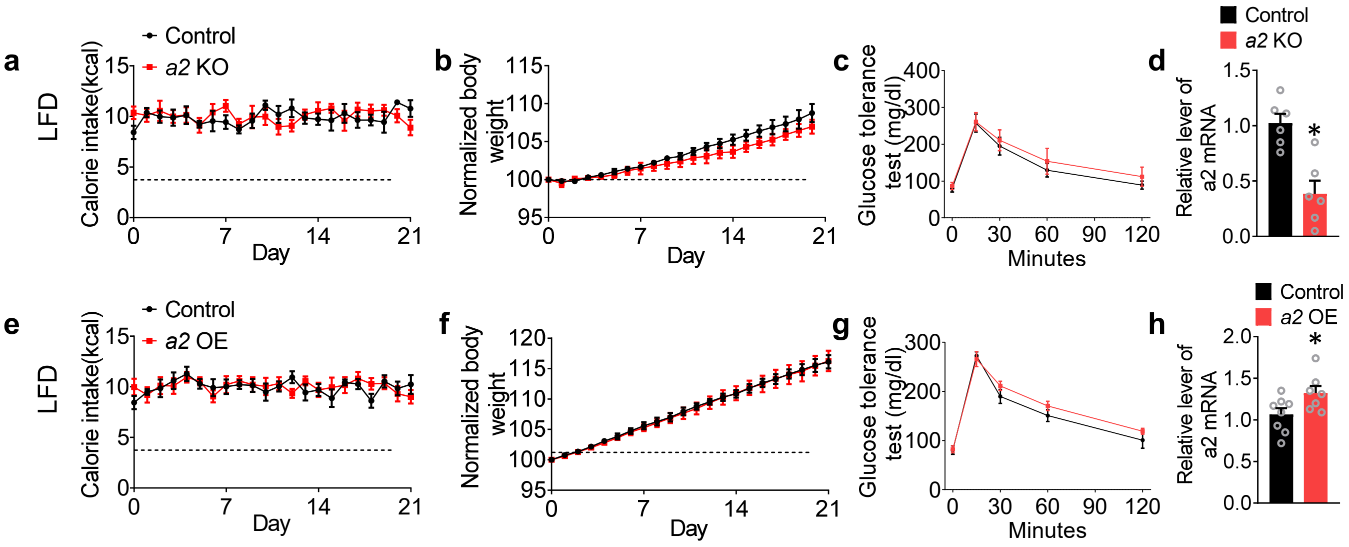


Extended Data Fig.4
