## Supplementary material for "Reversal of obesogenic feeding and hypometabolism by a bifurcating GABAergic neural circuit": Methods

**Materials and Methods**

Animals

All animal care and experimental procedures were approved by the Institutional Animal Care and Use Committees at Baylor College of Medicine. Mice used for data collection were at least eight weeks old males; and kept in temperature- and humidity-controlled rooms, in a 12/12 hr light/dark cycle, with lights on from 6:00 AM-6:00 PM. Health status was normal for all animals. Genetic Mouse Model. $Agrp^{DTR/+}$ mice[1], $Agrp^{Cre}$ mice[2], $Agrp^{nsCre}$ mice[3], $Lepr^{lox/lox}$ mice[4], Ai32 mice[5], $Npy^{GFP}$ mice[6], $Gad1^{lox/lox}$::$Gad2^{lox/lox}$ mice[3], $Rosa26^{tdTomato}$ mice[5], $Mc4r^{Cre}$ mice[7], and $Rosa26^{Cas9}$ mice[8] were produced as described previously. All mice are on a C57Bl/6 background with at least 8 generations backcrossed.

Stereotaxic Surgery

All the mice with brain surgery were performed with the same pre-operative and post-operative care as described previously[9]. Briefly, animals were received analgesia buprenorphine (s.c.) one hour prior to the start of anesthesia then anesthetized with isoflurane and placed on a stereotaxic frame (David Kopf, Tujunga, CA). The local anesthetics was applied before making an incision. All surgeries were performed on a heating pad and allowed to recover in a heating cage.

To perform the viral injection, virus was loaded into a needle (Hamilton Small Hub RN 33G, Reno, NV) connected with a 10 µL syringe (Hamilton 700 Microliter, Reno, NV). Injections were performed with an Ultra MicroPump (World Precision Instruments, Sarasota, FL) and Micro4 Controller (Heidenhain Corporation, Schaumburg, IL), at a rate of 0.1µL/min. For AAV a total 0.3 µl volume was delivered into brain regions. For HSV a total 0.2 µl was delivered into the DMH or MPO. The relevant stereotaxic coordinates for the injections are described in the following according to a standardized atlas of the mouse brain (Franklin and Paxinos, third edition, 2007).

For viral injection, the coordinate of ARC is AP = -2.06 mm, ML = ± 0.25 mm, DV = -5.9 mm; the coordinate of DMH is AP = -1.94 mm, ML = ± 0.3 mm, DV = -5.3 mm; and the coordinate of MPO is AP = -0.1 mm, ML = ± 0.2 mm, DV = -5.4 mm.

For cannulation of optical fiber, the coordinate of DMH is AP -1.94 mm, ML + 0.3 mm, DV -4.9 mm). The coordinate of DMH is AP -0.1 mm, ML + 0.2 mm, DV -5.0 mm.

The virus *AAV2-EF1a-DIO-hChR2(E123T/T159C)-GFP* was from UNC; the *HSV-LSL-EYFP* and *HSV-LSL-mCherry* were from Harvard; the plasmids *AAV9-fDIO-WGA-nsCre-mCitrine, AAV9-fDIO-WGA-mCitrine, AAV9-DIO-Gabra3*$^{sgRNA}$*-tdTomato*, *AAV9-DIO-Gabra3*$^{cDNA}$*-tdTomato, AAV9-DIO-Gabra2*$^{sgRNA}$*-tdTomato*, and *AAV9-DIO-Gabra2*$^{cDNA}$*-tdTomato* were synthesized by our lab. All noncommercial viruses including *AAV9-DIO-WGA-zsGreen*[10] were packaged by Optogenetics and Viral Design/Expression Core at Baylor College of Medicine. All of virus were diluted to a final working titer with no less than 2 x $10^{12}$ viral genomes per ml.

Ablation of AgRP Neurons in neonates

To ablate AgRP neurons, the newborn pups were either injected with DT (75 ng in 20 μl saline, s.c.) or saline. After weaning and applying ear tags, the genotype was determined by PCR. The 28 days after DT injection the mice were treated with leptin (4.0mg/kg, i.p. twice per day) for 3 days. The mice were euthanized, and the brain was harvested. The whole brain was sectioned with 20 μm thickness by a microtome (ThermoFisher Scientific, Waltham, MA). Immunostaining for Fos was performed. Fluorescent images of Cy2-labeled Fos+ neurons in brain regions were obtained by an Axio Observer microscope (Zeiss, Thornwood, NY) and further analyzed using ImageJ software (NIH).

Optogenetics

For *in vivo* optogenetic stimulation, the optic fiber was assembled as described following the protocol [11]. To perform the photostimulation the optical fiber was connected to spectralynx (Neuralynx, Inc, USA) through a patch cable. For the food intake measurement, the blue light was shed into the DMH or MPO at 20 Hz with 10 ms pulse for 1 hour. The power of laser (0.5-1.2 mW) was calculated by optical power meter (PM100D, Thorlabs, Newton, NJ) before each experiment.

For *in vitro* optogenetics, the optic fiber was assembled as described following the protocol [11]. The blue light was controlled by a pulse stimulator. The blue light pulses (20 Hz, 10 ms/pulse) were shed onto the ChR2-expressing AgRP axonal fibers within the DMH or MPO. The power of the laser (0.5-1.2 mW) was measured by a power meter (PM100D, Thorlabs, Newton, NJ) before experiments.

Drug Administration

To general administration of drugs, the mice were received i.p. injection of leptin at a dose of 4.0 mg/kg 30 min before feeding assay.

For infusing bicuculline (Bic) into the DMH or MPO the guide cannula (23-gague steel) was obtained from Plastics One (Roanoke, VA). A circular craniotomy (diameter 0.5 mm) was drilled at the locations of DMH or MPO. The guide cannula was installed on the holder and guided into the target brain region. To deliver drug the internal cannula was inserted onto the top of the guide cannula and extended below the guide cannula 0.5 mm. To target the DMH the cannula was implanted within the DMH with the coordinate (AP -1.94 mm, ML + 0.3 mm, DV -4.8 mm). To target the MPO the cannula was implanted within the MPO with the coordinate (AP -0.1 mm, ML + 0.2 mm, DV -4.9 mm). Bic (4 ng/side) was infused.

To disrupt GABAergic inputs from AgRP neurons in adult mice, microinjection of NB124[3] (two injections of 0.4 mg/side, 2 days apart; Calbiochem) in 8-week-old $Agrp^{nsCre/+}$::$Lepr^{lox/lox}$::$Jax2356^{NeoR/+}$::$Rosa26^{tdTomato}$ mice was performed.

Food Intake

For the acute feeding studies in optogenetic experiments, food intake was measured (from the start of the “lights off” cycle, 6 pm - 7 pm) one hour during and after 1-hr photostimulation with chow diet (5V5R, LabDiet, St. Louis, MO) or high-fat diet (HFD, 60% kcals from fat, Bio-Serv).

The regular food intake (4 hrs) in well-fed mice was monitored from 6 pm to 10 pm. The feeding test were performed 3 weeks after virus injection. For the chronic feeding studies, food intake as well as body weight was daily measured between 9 am and 10 am up to 4 weeks.

Glucose Tolerance Test (GTT)

The GTT was performed as described[3]. Briefly, mice were fasted overnight for 16 h, a blood sample was taken, and then the mice were injected with D-glucose (1 g/kg, i.p.), and blood was drawn from the tail vein at 0, 15, 30, 60, and 120 min later. Blood glucose levels were determined with a FreeStyle Lite glucometer (Abbott Laboratories).

Energy Expenditure

$O_2$ consumption, $CO_2$ production, and RQ were monitored by Comprehensive Lab Animal Monitoring System (CLAMS; Columbus Instruments, Columbus, OH) [3]. Mice were acclimatized in the chambers for 48 hrs prior to data collection.

*In vitro* Electrophysiology

The brains of adult mice were sectioned in coronal plane (250 − 300 μm). The brains and slices were handled and kept in artificial cerebrospinal fluid (aCSF) as described recently [12]. Animals were subjected to anesthesia, and the handling protocol of the local committee was followed. In most case we have used the same mouse first for the *in vivo* experiments and immediately after for combined electrophysiology and optogenetics *in vitro* in order to minimize the number of mice. Mice were deeply anesthetized with isoflurane and transcardially perfused with a modified ice-cold sucrose-based cutting solution (pH 7.3) containing 10 mM NaCl, 25 mM $NaHCO_3$, 195 mM Sucrose, 5 mM Glucose, 2.5 mM KCl, 1.25 mM $NaH_2PO_4$, 2 mM Na-Pyruvate, 0.5 mM $CaCl_2$, and 7 mM $MgCl_2$, bubbled continuously with 95% $O_2$ and 5% $CO_2$. The mice were then decapitated, and the entire brain was removed and immediately submerged in the cutting solution. Slices were cut with a Microm HM 650V vibratome (Thermo Scientific). Slices containing the DMH were recovered for 1 h at 34°C and then maintained at room temperature in artificial cerebrospinal fluid (aCSF, pH 7.3) containing 126 mM NaCl, 2.5 mM KCl, 2.4 mM $CaCl_2$, 1.2 mM $NaH_2PO_4$, 1.2 mM $MgCl_2$, 11.1 mM glucose, and 21.4 mM $NaHCO_3$) saturated with 95% $O_2$ and 5% $CO_2$ before recording. Slices were transferred to a recording chamber and allowed to equilibrate for at least 10 min before recording. The slices were superfused at 34°C in oxygenated aCSF at a flow rate of 1.8-2 ml/min. ZsGreen-labeled neurons in the DMH were visualized using epifluorescence and IR-DIC imaging on an upright microscope (Eclipse FN-1, Nikon) equipped with a moveable stage (MP-285, Sutter Instrument). Patch pipettes with resistances of 3-5 MΩ were filled with intracellular solution (pH 7.3) containing 128 mM K-Gluconate, 10 mM KCl, 10 mM HEPES, 0.1 mM EGTA, 2 mM MgCl2, 0.05 mM Na-GTP and 0.05 mM Mg-ATP. Recordings were made using a MultiClamp 700B amplifier (Axon Instrument), sampled using Digidata 1440A and analyzed offline with pClamp 10.3 software (Axon Instruments). Series resistance was monitored during the recording, and the values were generally <10 MΩ and were not compensated. The liquid junction potential was +12.5 mV and was corrected after the experiment. Data were excluded if the series resistance increased dramatically during the experiment or without overshoot for action potential. Currents were amplified, filtered at 1 kHz, and digitized at 20 kHz. Current clamp was engaged to test neural firing frequency and resting membrane potential (Vm) at the baseline. The aCSF solution contained 1 μM tetrodotoxin (TTX) and a cocktail of fast synaptic inhibitors, AP-5 (30 μM; an NMDA receptor antagonist) and DNQX (30 μM; an AMPA receptor antagonist) to block the majority of presynaptic inputs. For the light evoked inhibitory postsynaptic current (IPSC) recordings, the internal recording solution contained: 125 mM $CsCH_3SO_3$; 10 mM CsCl; 5 mM NaCl; 2 mM $MgCl_2$; 1 mM EGTA; 10 mM HEPES; 5 mM (Mg)ATP; 0.3 mM (Na)GTP (pH 7.3 with NaOH). IPSC within the DMH neurons was measured in the current clamp mode in the presence of 1 μM TTX and 4-AP with or without 50 μM bicuculline.

*In vivo* Tetrode Recording

We used the microdrive model that enabled deliver laser or drug into brain and recording neural activities simultaneously [13,14]. The microdrives were modified on the basis of tetrode microdrives from Neuralynx which were loaded with one optic fiber in the centre and 7 nichrome tetrodes consisting of 4 thin wires twined together (STABLOHM 675, California Fine Wire Co., Grover Beach, CA). The optic fiber positioning 0.1 mm from the tetrode was glued to the middle of the bundle of tetrodes. Tetrode tips were goldplated to reduce impedance to 0.3– 0.4 M (tested at 1 kHz). The microdrive was implanted into the DMH or MPO in the *Mc4r^Cre^* mice with injection of *AAV2-EF1a-DIO-hChR2(E123T/T159C)-GFP* within the DMH or MPO. After recovery from microdrive implantation, the mouse was connected to a 32-channel preamplifier headstage. All signals recorded from each tetrode were amplified, filtered between 0.3 kHz and 6 kHz, and digitized at 32 kHz. The local field potentials were amplified and filtered between 0.1 Hz and 1 kHz. The tetrodes were slowly lowered in quarter-turns of a screw on the microdrive (~60-μm steps). Spikes were sorted using Offline Sorter software (Plexon). Units were separated by the T-Distribution E-M method, and cross-correlation and autocorrelation analyses were used to confirm unit separation. Clustered waveforms were subsequently analyzed by using NeuroExplorer (Nex Technologies, Colorado Springs, CO) or MATLAB (MathWorks, Natick, MA). The firing rates were presented with spikes per bin with 1 s interval or spikes per second. The ChR2+ neurons were identified by the short latencies of evoked spikes accurately following high-frequency photostimulation, as well as the identical waveforms of evoked and spontaneous spikes [15].

Real-Time qPCR.

Sorted cells were immediately lysed by RLT buffer from the RNeasy Plus Micro kit (Qiagen). Total mRNA was subsequently extracted and purified based on the manufacturer's suggested protocol (Qiagen). For some experiments, total RNA was extracted from fresh arcuate nucleus by TRIzol (Invitrogen) per the manufacturer's instructions. The purified RNA was quantified by One-drop spectrophotometer (Thermo Fisher), and mRNA was reverse-transcribed by using the SuperScript II kit (Invitrogen) per the manufacturer's suggested protocol. qPCR was performed in the Bio-Rad CFX96 Real-Time PCR system with a set of Taqman probes (IDT). Relative abundance of *Lepr*, *α3*, or *α2* transcripts was determined by using the $2^{-\Delta\Delta Ct}$ method and normalized to Gapdh, a housekeeping gene.

Histology

Immunostaining was performed as described with modification [3]. Mice were killed and perfused transcardially with ice-cold PBS buffer (pH 7.4) containing 3% (wt/vol) paraformaldehyde (Alfa Aesar) and 1% glutaraldehyde (Sigma, St. Louis, MO). Brains were collected and postfixed overnight under 4 °C in a fixation buffer containing 3% paraformaldehyde. Free-floating sections (25 μm) were cut by a microtome (ThermoFisher Scientific, Waltham, MA) and then blocked with 5% (wt/vol) normal donkey serum in 0.1% Triton X-100 (TBST buffer, pH 7.2) for overnight. Goat anti-AgRP (1:500 dilution; Santa Cruz Biotech, Dallas, TX) or rabbit anti-Fos (1:1,500 dilution; EMD Millipore, Burlington, MA) was applied to the sections for overnight incubation under 4 °C, followed by 4 × 15-min rinses in the TBST buffer. Finally, sections were incubated with Alex Fluor Cy2-conjugated secondary antibody (1:1,000 dilution; Jackson Immunolab, West Grove, PA) for 2 h at room temperature, followed by 4 × 15-min rinses in TBST buffer. For mounted sections, fluorescent images were captured by a digital camera mounted on an Axio Observer microscope (Zeiss, Thornwood, NY).

Fluorescent in situ hybridization (FISH) was performed to examine the expression of Mc4r in the DMH and MPO. All reagents and Mc4r probe are commercially available from Advanced Cell Diagnostics (Newark, CA). The RNAscope Fluorescent Assay is one of FISH technique to visualize cellular RNA targets in fresh frozen tissues. All FISH procedures were performed following the protocol provided by the manufacturer. Briefly, fresh frozen brain tissues were sectioned with cryostat at 20 μm thickness and mounted onto SuperFrost Plus slides. Chill slides into the 4% PFA to fix for 15 min at 4°C. Then the sections were dehydrated with grade ethanol. Air dry slides for 5 min at RT, add ~5 drops of RNAscope Protease IV to each section for 30 min at RT and wash 3 times. The sections were hybridized Drd1 probe for 2 hrs at 40°C. After 2 times washes, the slides were hybridized Amp1FL for 30 min, Amp-2FL for 15 min, Amp-3FL for 30 min, and Amp-4FL for 15 min at 40°C followed by washing 3 times. Finally, coverslip over sections. The fluorescent images were captured by Axio Observer microscope.

Statistical Analyses

Data were analyzed by unpaired t test, paired t test, one-way or two-way ANOVA with the post hoc as appropriate. Wilcoxon signed rank test was used when the data were not normally distributed. Statistical analyses were performed using Prism software (GraphPad Software, San Diego, CA) Results were considered significantly different at $p < 0.05$. All data are presented as mean ± S.E.M.
